# Direct Detection and Atomic Modeling of Ligands in Cryo-EM Maps Using Deep Learning

**DOI:** 10.64898/2026.06.01.729423

**Authors:** Shu Li, Anika Jain, Yuki Kagaya, Joon Hong Park, Daisuke Kihara

**Affiliations:** Department of Computer Science, Purdue University, West Lafayette, IN, United States; Department of Biological Sciences, Purdue University, West Lafayette, IN, United States

## Abstract

Cryogenic electron microscopy (cryo-EM) has become an increasingly important for structure-based drug discovery by enabling characterization of interactions between macromolecules and small-molecule ligands. However, computational interpretation of ligand density remains challenging, particularly when ligand locations are unknown or local map resolution is limited. Existing methods generally require well-resolved macromolecular structures and predefined binding sites, limiting their applicability during early-stage structure determination. To date, no approach has been able to both reliably detect ligand density and subsequently reconstruct ligand atomic structures directly from experimental cryo-EM maps.

Here, we present Emap2lig, a two-stage deep learning framework for automated ligand detection and atomic modeling directly from cryo-EM maps. Emap2lig-Find identifies ligand-associated densities and remains effective for maps at resolutions as low as ∼5 Å. Emap2lig-Build subsequently uses a diffusion-based generative model to build atomic ligand structures. Together, Emap2lig provides a unified framework for ligand discovery and modeling across a broad range of resolutions.

## Introduction

Cryo-electron microscopy (cryo-EM) has emerged as a popular method for resolving the structures of biomolecules. With the rapid improvement in resolution of cryo-EM structures due to more efficient electron detectors and image processing algorithms, cryo-EM is fast gaining significance for not just obtaining macromolecular structures but also studying the interactions of these structures with small molecule ligands to aid in Structure-based drug discovery (SBDD) ^1^. The structures of many important protein-small molecule targets have been obtained using cryo-EM, such as allosteric modulators of G-protein coupled receptor GLP-1R that revealed important insights into conformational changes upon ligand binding that were not apparent in X-ray structures^2^.

Many computational methods have been developed to support each stage of cryo-EM data processing from 2D micrographs to three-dimensional map reconstruction, and macromolecular structure building and refinement^3^. In recent years, deep learning has been successfully applied across multiple tasks, such as 2D particle picking^4,5^, conformational analysis^6^, macromolecular structure modeling^7–9^, and model validation^3,10^. Several approaches have also been proposed for modeling ligand conformations in cryo-EM density maps. GLIDE-EM^11^ extends GLIDE molecular docking by incorporating cryo-EM density information. Chem-EM^12^ employs density-guided docking strategies that require an accurately modeled protein structure and a predefined binding pocket, followed by local refinement. EMERALD^13^ infers ligand structures by converting ligand density into a pseudo-atomic skeleton. These methods are designed for cases in which the macromolecular structure is already well resolved, and the ligand-binding pocket is clearly defined in a high-quality map. However, such approaches are less suitable at earlier stages of structure determination, when ligand positions are unknown, binding sites are unexpected, or ligand density is locally weak or poorly resolved. To the best of our knowledge, no existing method can directly detect ligand densities within a cryo-EM map of a biomolecular complex without prior knowledge of ligand location.

Here, we present Emap2lig, a two-stage deep learning framework for ligand detection and modeling directly from experimental cryo-EM density maps. Emap2lig consists of Emap2lig-Find, which identifies ligand-like densities within a map, and Emap2lig-Build, which models the atomic coordinates of ligands within the detected densities when the ligand identity is known. A key advantage of Emap2lig is that it requires no prior knowledge of ligand locations or binding pockets. Emap2lig-Find employs a U-net–based neural network to detect all potential ligand densities in a cryo-EM map. Subsequently, Emap2lig-Build performs instance segmentation to isolate individual ligand densities and guides a structure module to generate three-dimensional atomic models of the ligands. This design enables Emap2lig to operate directly on complex maps containing full biomolecular assemblies, providing a unified solution for ligand discovery from cryo-EM data and facilitating structure-based drug discovery. Emap2lig was designed to detect and model ligands at resolutions up to ∼5 Å. In the resolution range of 0–3 Å, it achieved a map-level recall of 0.73, while in the 3–6 Å range the recall was 0.54. Unlike existing approaches, Emap2lig-Build can successfully model large ligands containing up to 80 atoms in near native poses Ligand conformations built by Emap2lig-Build has a mean structural deviation of 1.57 Å from native structures. Emap2lig is available as a webserver for a convenient use (https://em.kiharalab.org/algorithm/Emap2lig-Find and https://em.kiharalab.org/algorithm/Emap2lig-Build).

### Workflow of Emap2lig

In **Fig. 1** we overview the network architecture of Emap2lig. The architecture of Emap2lig is composed of two distinct functional stages, Emap2lig-Find for ligand detection and Emap2lig-Build for precise structure modeling. The pipeline begins with Emap2lig-Find, a module designed to detect ligand instances within the cryo-EM map without requiring prior knowledge of the ligand identity. The primary function of this stage is to perform an initial density segmentation of the full volume to locate potential binding sites. Utilizing a U-net model^14^ trained on 64 Å³ boxes (**Supplementary Fig. 1**), this module (blue box in **Fig. 1a**) classifies each voxel into one of six primary structural classes: backbone, sidechain of proteins, phosphate, sugar, base of nucleic acids, or ligand. This broad classification step effectively isolates ligand densities from the surrounding macromolecular structure’s densities, setting the stage for targeted modeling.

**Fig. 1.**
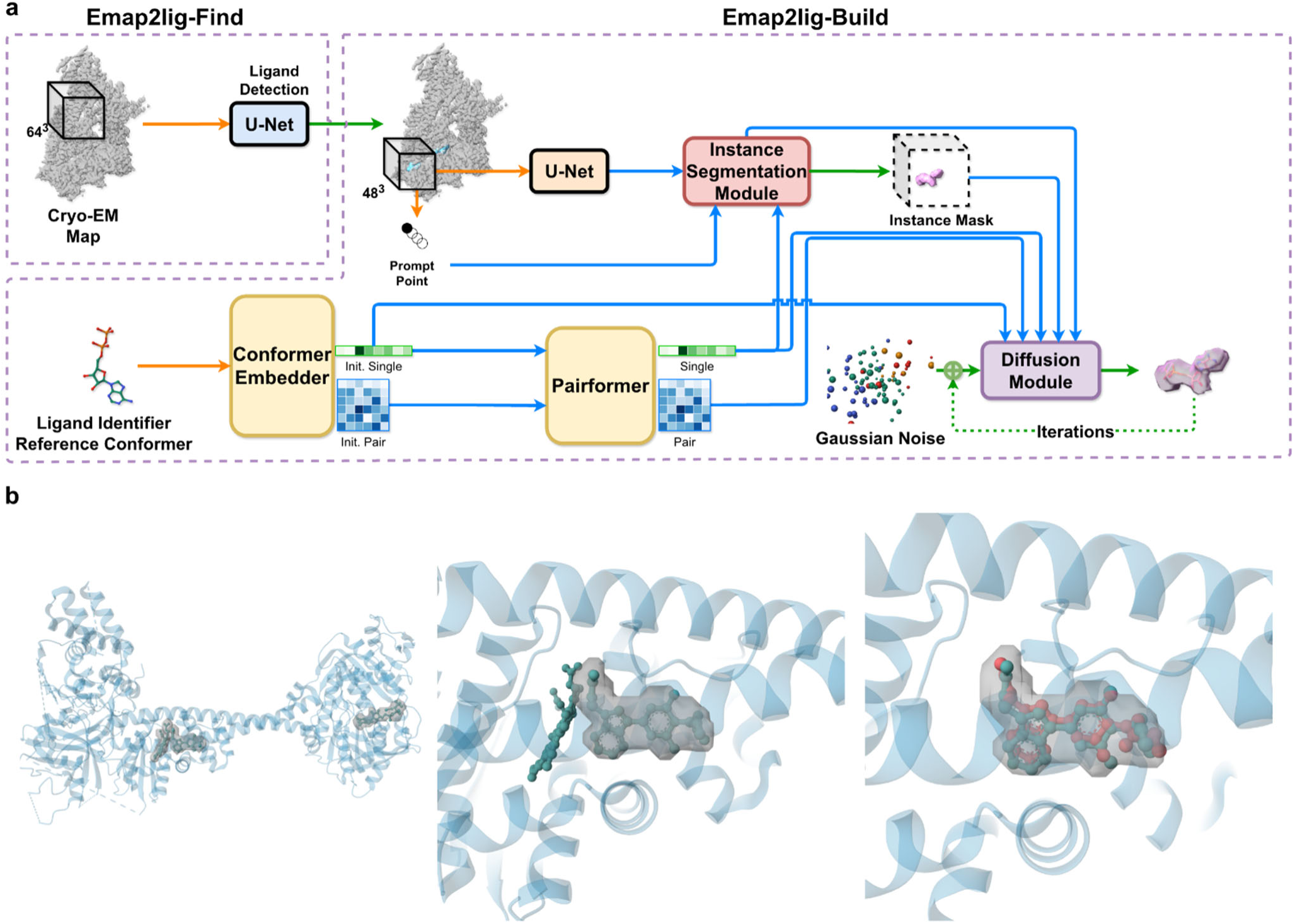
Network architecture of Emap2lig. **a**, Overall architecture of Emap2lig, which comprises Emap2lig-Find and Emap2lig-Build. Emap2lig-Find segments the input cryo-EM density map using a U-net to identify candidate ligand-binding regions within the density map. Emap2lig-Build generates a 3D ligand structure conditioned on a prompt point sampled from the detected region and on a reference conformer that provides prior chemical and geometric constraints. Within Emap2lig-Build, the conformer embedder encodes the reference conformer into initial single-atom and initial pairwise representations, and the pairformer updates and integrates these representations to propagate chemical constraints across the ligand atoms. The instance segmentation module refines a ligand-specific mask by integrating U-net voxel features with single-atom representations derived from pairformer. The refined mask defines the spatial region for structure generation, from which 8,192 points are sampled, while global features are extracted via adaptive 3D pooling to capture overall spatial context. The diffusion module then iteratively constructs the final 3D ligand structure through denoising, conditioned on initial single-atom representation from the conformer embedder, the single-atom and pairwise representations as structural constraints and on the point-wise feature and global feature from the instance segmentation module as spatial constraints. Orange arrows indicate data flow, blue arrows indicate intermediate module outputs, and green arrows indicate the final output for training loss calculation. **B**, An example of ligand detection and modeling with a cryo-EM map of human soluble guanylate cyclase with GZO (a PDB code), HEM, and G2P as bound ligands. (EMD-30618; resolution 3.7 Å. PDB: 7D9R). Left: identified ligand density (gray) generated by the Emap2lig-Find stage. Middle, the instance mask of GZO produced by the instance segmentation module. Notably, the adjacent HEM region is not included in the mask. Right: the ligand structure predicted by the diffusion module. The native macromolecular structure is depicted as a blue cartoon. The ligand structure in the PDB file is represented in blue using a ball-and-stick model, while the modelled ligand structure is shown in red using a ball-and-stick model.

The second stage, Emap2lig-Build, reconstructs the 3D structure of a ligand when its chemical identity is provided by users. This stage is trained as a unified framework and comprises four functional modules that sequentially integrate cryo-EM density and chemical information. The first module of the Build stage utilizes a distinct U-net (the orange box in **Fig. 1a**) to reprocess local density, identifying features explicitly tailored for structure generation. Running concurrently with volumetric cryo-EM feature extraction, the conformer embedder and the pairformer modules encode spatially invariant chemical priors derived from a computationally generated reference conformer of the ligand provided by users. This reference geometry is initialized via RDKit^15^ utilizing the ETKDGv3 (Experimental-Torsion-Knowledge Distance Geometry) algorithm^16^ to obtain a low-energy 3D conformation. Although the global topology of this reference conformer, particularly flexible dihedral angles, may differ from the bound bioactive conformation, its local geometric properties, such as covalent bond lengths, bond angles, and stereochemistry, remain physically consistent. The conformer embedder converts this reference conformer into initial single-atom and pairwise representations, and the pairformer propagates these constraints and updates single and pairwise representations. By incorporating these chemically grounded priors, the model recovers bond-level information that is often ambiguous or obscured in noisy cryo-EM density maps.

The subsequent step, the instance segmentation module (**Supplementary Fig. 2**), is responsible for extracting the precise mask for the target ligand. It integrates the voxel features from the U-net (blue arrow in **Fig. 1a**) with the single representation from the pairformer. Finally, the diffusion module (**Supplementary Fig. 3**) refines ligand coordinates to produce the final 3D conformer (**Fig. 1a**). Five contextual inputs guide this process: the initial 1D single-atom representation from the conformer embedder, the 1D single-atom representation and 2D pairwise representation from the pairformer, which encode atom-level features and enforce stereochemical bonding constraints, as well as the point-wise features and global features from the instance segmentation module, which define spatial boundaries within the map. By jointly conditioning on chemical priors and spatial constraints, the diffusion trajectory converges toward a conformation that is consistent with the experimental density while maintaining chemical validity.

The model was trained using a rigorously curated dataset sourced from the Electron Microscopy Data Bank (EMDB)^17^ and Protein Data Bank (PDB)^18^, comprising 7,471 high-quality cryo-EM maps with resolutions ranging from 1.4 to 6.0 Å. To prevent homologous structure leakage and ensure robust evaluation, we performed sequence clustering to partition the macromolecular targets into structurally non-redundant subsets. This yielded a dataset of 94,379 ligand instances (representing 16,359 unique ligand types), partitioned into a training set (7,065 maps; 91,814 instances), a validation set (178 maps; 1,171 instances), and a test set (228 maps; 1,394 instances). The training and validation dataset entries are provided in **Supplementary Table S1,** and the test dataset entries are provided in **Supplementary Table S2**. The training proceeded in three stages: following the initial training of the Emap2lig-Find detection module, the conformer embedder, pairformer, and instance segmentation modules were jointly pre-trained on the full dataset. Finally, we trained the Emap2lig-Build components on a high-confidence subset of 72,938 ligand instances, filtered using a segmentation recall threshold of 0.3. This filtering step ensured that the generative diffusion process was conditioned on reliable volumetric signals. Additional details are provided in the Methods section.

### Evaluation of Emap2lig-Find

We first evaluated Emap2lig-Find, the ligand detection module for EM maps (**Fig. 1**). During evaluation, predictions were binarized into predicted ligand/non-ligand voxels using Li’s minimum cross-entropy threshold^19^ computed independently on the validation set. **Fig. 2a** shows the overall detection performance evaluated at both the map level and the individual ligand–instance level. Map-level detection accuracy was assessed using three metrics, HitRate at 30% and 50% coverage (HR30 and HR50) as well as overall recall (three plots in blue). HR30 and HR50 are defined as the fraction of ligands in a map for which at least 30% or 50% of the grid points were correctly identified as ligand density:

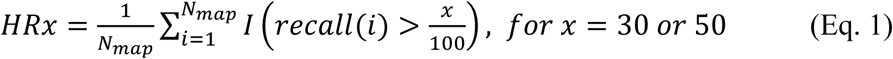

**Fig. 2.**
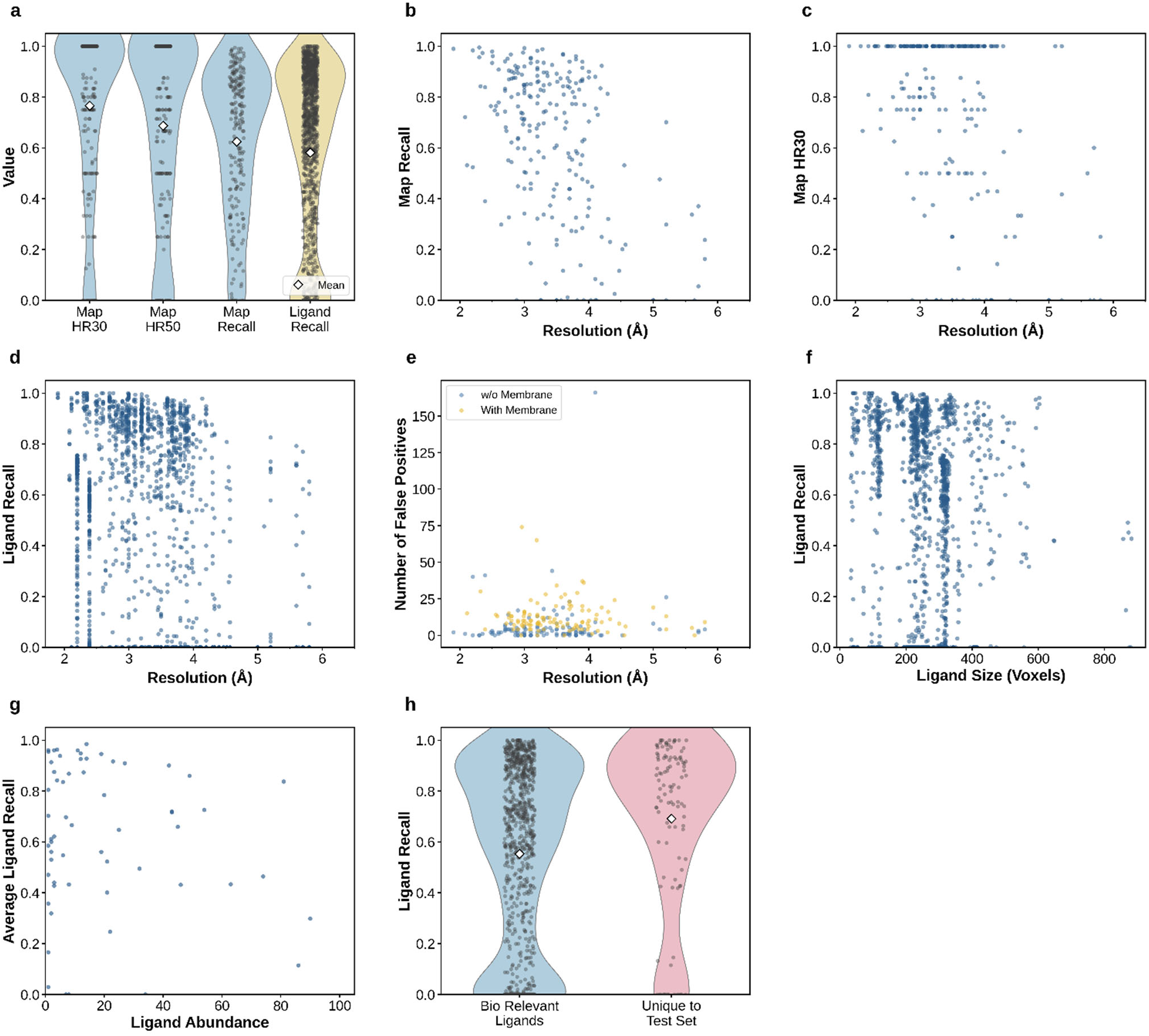
Evaluation of Emap2lig-Find. **a.** Distribution of map-level and ligand instance-level detection metrics. White diamonds indicate means. Three map-level metrics are evaluated over 228 maps in the test set. HR30, the fraction of ligands per map detected with at least 30% voxel recall (median:1.00, mean: 0.77); HR50, the fraction detected with at least 50% voxel recall (median: 0.80, mean: 0.69); and map recall, the average per-ligand recall within each map (median: 0.72, mean: 0.62). The instance-level metric reports the per-ligand recall for each of 1,394 individual ligand instances in the test set (median: 0.70, mean: 0.58), defined as the fraction of the native ligand voxels correctly identified. **b.** Map-level recall relative to the map resolution (Å) for the 228 test maps (mean resolution: 3.4 Å). **c.** Map HR30 relative to the map resolution (Å) for the same 228 test maps. **d.** Instance-level ligand recall relative to the map resolution (Å) for 1,394 ligand instances in the test set. **e.** The number of false-positive predicted ligand density per map relative to the map resolution, stratified by the presence of membrane stabilization systems (e.g., nanodiscs). A false-positive prediction is a predicted ligand density region that does not overlap with any ligand in the reference PDB file. Maps containing membrane stabilization systems (n: 108 maps; mean: 12.4, median: 9.5 false positives per map) produced substantially more false positives than maps without them (n: 120 maps; mean: 6.3, median: 2 false positives per map). **f.** Ligand recall relative to the ligand size measured as voxel counts (mean ligand size: 253 voxels). **g.** The Average ligand recall in the test set per ligand type relative to its occurrence counts in the training set, shown for ligand types. 8 ligands with more than 100 counts are omitted in this plot. **h.** Comparison of ligand-level recall between all biologically relevant ligands in the test set (908 ligands spanning 68 unique ligand types; median recall: 0.66, mean: 0.55) and the subset of them whose chemical identity was absent from the training set (110 ligands; spanning 26 unique ligand types; median recall: 0.82, mean: 0.69).

where *N_map_* is the number of ligands in the map, recall for a ligand *i* is defined as the proportion of correctly identified grid points as ligand density among all the grids that belong to the ligand *i* in the EM map. *I*(⋅) is the indicator function that equals 1 if the condition is true and 0 otherwise. The overall recall evaluates the fraction of correctly identified grid points that belong to ligand density in an entire map.

The median HR30 and HR50 values were 1.00 and 0.80, respectively (**Fig. 2a**), indicating that Emap2lig-Find identified ligand density for most ligands in each map. Notably, all ligands were detected in 198 (86.8%) out of 228 maps at the 30% (HR30) and 172 maps (75.4%) at the 50% (HR50) thresholds, respectively. The median map-level recall was 0.72, indicating that 72% of ligand density was correctly detected. At the individual ligand level (yellow plot), the median recall is 0.70. Overall, these results indicate that Emap2lig-Find can successfully identify ligands in EM maps, despite the very small fraction of the map volume (0.83%) occupied by ligand density.

Ligand detection is only weakly influenced by map resolution when evaluated over all ligand voxels at the map level (**Fig. 2b**). This dependence is further reduced when using the HR30 metric, indicating that most ligands were detected sufficiently with little influence from map resolution (**Fig. 2c**). Results using HR50 are provided in **Supplementary Fig. 4**, which shows the same trend. **Fig. 2d** further illustrates the effect of map resolution at the individual ligand level. The majority of ligands were detected with high recall, with 50.7% of ligands achieving a recall greater than 0.7. Individual results are provided in **Supplementary Table S3.**

We observed that 15 of 228 maps (6.7%) had zero recall at the map level (**Fig. 2b**). Among these, four maps had resolutions worse than 5 Å, where ligand density appeared as continuous with surrounding macromolecular density, making it difficult to identify ligand-like features. Nine maps contained ligands that were mispredicted as proteins or nucleotides. In most instances, the ligand density at the author-recommended contour level lacked a distinct shape. In EMD-27416, U5P (uridine-5′-monophosphate) was located in close proximity to the terminal nucleotide of the chain, which may have led to its misclassification as an extension of the nucleotide chain rather than a standalone ligand. Finally, two maps contained all ligands at the interface with membrane stabilization systems. These regions exhibit noisy density due to contributions from lipid chains, proteins, and detergent molecules. A further challenge is that PDB does not consistently annotate all lipid molecules, despite all being technically ligands, making it difficult to train models that can reliably distinguish biologically relevant ligands from components of membrane stabilization systems.

In **Fig. 2e** we conversely analyzed the number of false positives as a function of map resolution in the subsequent plot. Maps of biological systems were separated into those containing membrane components (yellow) and those without membranes (blue), because membrane regions contain large numbers of lipids, which are technically treated as ligands in Emap2lig. The presence of membrane components complicates the evaluation of ligand detection accuracy, since most lipids in membrane regions are not annotated in PDB files. As shown in **Fig. 2e**, maps with membrane components exhibit substantially more apparent false positives than maps without membranes (individual data is provided in **Supplementary Table S4**). The median number of false positives is ten for maps with membranes, compared with two for maps without membranes. No clear dependence on map resolution is observed. Related to **Fig. 2e**, map level recall relative to map resolution is provided in **Supplementary Fig. 5**. In terms of recall, membrane systems did not affect negatively.

The remaining three panels further examine ligand-level recall with respect to ligand properties and types. **Fig. 2f** evaluates recall as a function of ligand size, defined by the number of voxels in the ligand density (**Supplementary Table S5**). Although small ligands might be expected to be more difficult to detect, no such trend is observed. Next, in **Fig. 2g**, we assess whether ligand detection is influenced by the frequency of ligand types in the training set (**Supplementary Table S6**). Despite the expectation that rare ligands (i.e., those with low abundance in the training data) would be more challenging to detect, no such trend is observed. Unseen ligands were identified using ligand similarity using SIMCOMP^20^ with complete linkage. Using a similarity threshold of 0.47, which resulted in 933 clusters, ligands in the test set that did not cluster with any training-set ligands were considered sufficiently dissimilar to be treated as unseen. Based on this clustering, we identified 32 unseen ligands in the test set. In **Fig. 2h**, ligand recall is compared between ligand types that were present in the training set and those that were entirely unseen during training (individual data in **Supplementary Table S7**). Notably, unseen ligands do not exhibit reduced recall, further supporting the observation from **Fig. 2g** that rare or even unseen ligands are detected with comparable accuracy to more common ligand types.

### Examples of detected ligands

**Fig. 3** shows eight examples of ligand density detected by Emap2lig-Find. We start with successful examples where ligand density was correctly identified with no false positive regions. In the first example (**Fig. 3a**) of a viral protein structure resolved at 3.3 Å (EMD-13641), two flavin adenine dinucleotide (FAD) molecules were correctly detected, achieving a high map-level recall of 0.93. **Fig. 3b** shows a lower-resolution (4.2 Å) map of an apoptosis-related protein hexamer (EMD-36450). All six ATP molecules were clearly detected, resulting in a recall of 0.92. **Fig. 3c** presents a membrane-associated structure at 3.75 Å resolution (EMD-34941). The presence of membrane density often increases the difficulty of ligand detection, as discussed above. In this map of an ABC transporter, two ATP molecules were correctly identified (recall: 0.82). Three false-positive regions were also predicted; however, they did not interfere with the correct ligand positions and could be readily distinguished as noise.

**Fig. 3.**
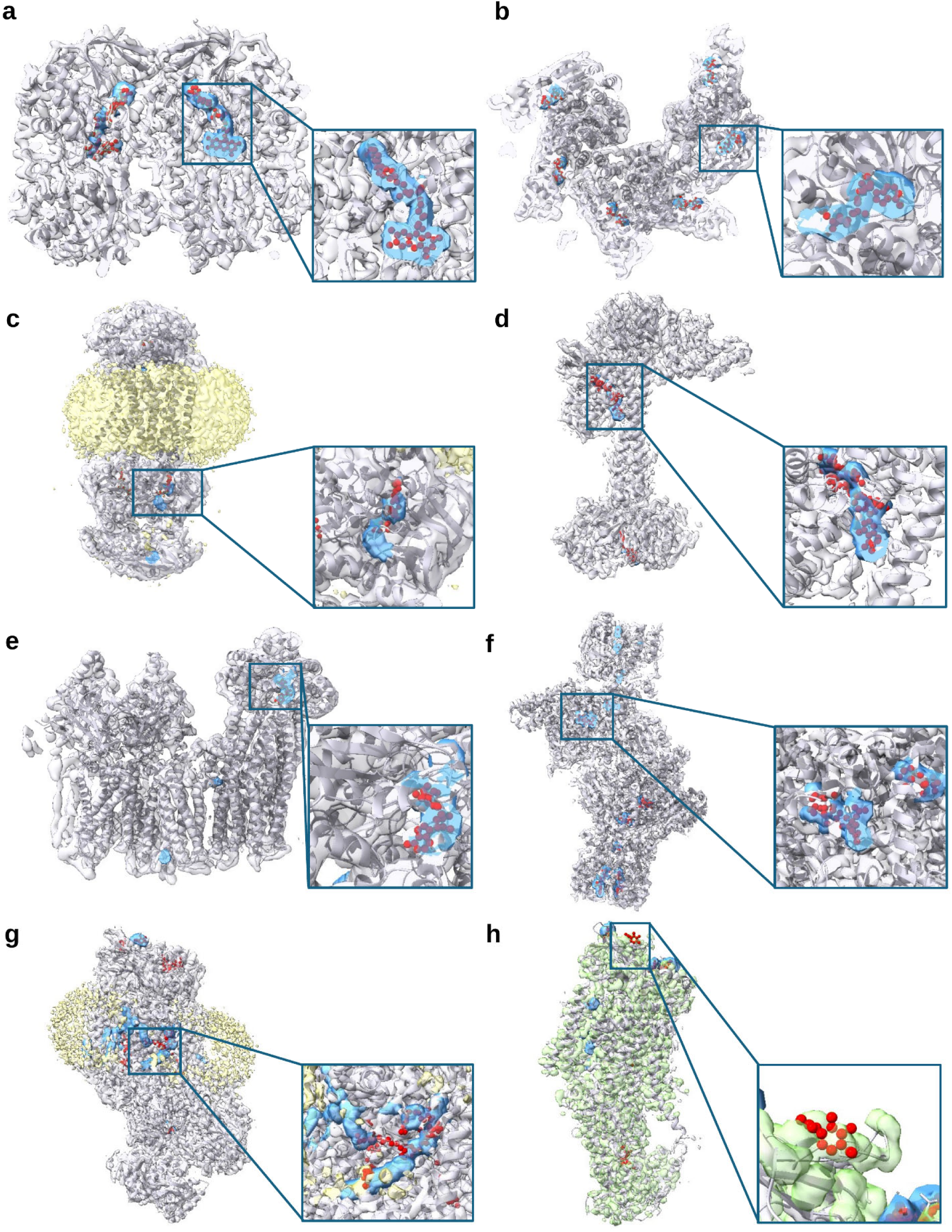
Examples of detected ligands in EM maps. The gray density represents the original EMDB map, and the gray cartoon corresponds to the associated deposited PDB structure. The blue density indicates ligand density detected by Emap2lig-Find, while the molecules in red are ligands in the PDB file. In panels c and g, yellow density represents the membrane stabilization system. **a**, Mimivirus genomic fibre assymetric unit (PDB: 7PTV; EMD-13641; resolution: 3.3 Å). The ligand shown in the inset is FAD. Map recall: 0.93. **b**, CED-4 hexamer (PDB: 8JNS; EMD-36450 res.: 4.2 Å). The ligand in the inset is ATP. Map recall: 0.92. **c**, Type I ABC transporter, LpqY-SugABC (PDB: 8HPR; EMD-34941; res.: 3.75 Å). Ligand in the inset is ATP. Map recall: 0.82. **d**, human soluble guanylate cyclase in NO+Rio state (PDB: 8HBF; EMD-34627; res.: 3.1 Å). Inset: GZO. Map recall: 0.93. **e**, pancreatic ATP-sensitive potassium channel (PDB: 7U7M; EMD-26309; res.: 5.2 Å). Inset: ATP. Map recall: 0.7. **f**, iron nitrogenase complex from *Rhodobacter capsulatus* (PDB: 8OIE; EMD-16890; res: 2.35 Å). Inset: S5Q. Map recall: 0.97. **g**, P4-ATPase(ATP8B1)-CDC50A complex in E2P active conformation with bound phosphatidylcholine (PDB: 8OX9; EMD-17261; res.: 2.72 Å). Inset: POV. Map recall: 0.67. **h,** ATP8B1-CDC50A in E1P-ADP conformation (PDB: 8OX5; EMD-17257; res.: 2.9 Å). Inset: NAG. The green density represents predicted sidechain density by Emap2lig-Find. Map recall: 0.44.

Critically, the method also demonstrated strong generalizability to structurally novel ligands (**Fig. 3d**). In a map of a pancreatic potassium channel (EMD-34627; 3.1 Å resolution), a drug molecule (PDB: GZO) that is dissimilar to any ligands in the training or validation sets (SIMCOMP^20^ score < 0.47) was successfully detected, achieving an instance recall of 0.93. Additionally, Phosphomethylphosphonic acid guanylate ester (G2P) at the bottom of the structure was fully detected with a ligand recall of 0.91.

Notably, the method remained effective even at resolutions worse than 5 Å. EMD-26309 (5.2 Å resolution) represents a particularly challenging regime for ligand detection, Emap2lig-Find was still able to identify ATP density with a map-level recall of 0.70 (**Fig. 3e**). The method further proved capable of handling maps with substantial ligand heterogeneity. A single map (EMD-16890; 2.35 Å resolution) contains six distinct ligands, ranging from a minimal four-heavy-atom ligand (AF3, aluminum fluoride) to complex iron–sulfur clusters (S5Q, FeFe cofactor; CLF, Fe₈S₇ cluster; and SF4, iron–sulfur cluster). Densities for all ligands were successfully detected, achieving a map-level recall of 0.97 (**Fig. 3f**).

In contrast, **Fig. 3g and 3h** illustrate cases where detected ligand densities do not fully agree with the annotated ligand molecules. **Fig. 3g** is a case of ligands that are located at the boundary of membrane and proteins. In a map of ATPase (ATP8B1)-CDC50A complex (EMD-17261; 2.72 Å resolution), eight false-positive regions were observed, all located within the membrane stabilization density. These predicted ligand densities exhibit elongated, lipid-like morphologies, suggesting that they may correspond to unannotated lipid densities rather than true false positives. The overall recall of 0.67 for the annotated ligand, POV, is largely attributable to incomplete detection of ligands located at the interface between the protein and the membrane environment. The final example (**Fig. 3h**) involves an N-acetylglucosamine (NAG) ligand (EMD-17257; 2.9 Å resolution). One NAG molecule was not detected because the model extended the asparagine side-chain classification into the adjacent ligand density. This behavior is plausible, as NAG is typically covalently linked to asparagine residues, resulting in a blurred boundary between the ligand and the protein side chain. Despite this limitation, the method successfully identified another isolated NAG ligand as well as a glycan chain (NAG–NAG–BMA).

### Detecting unmodelled ligands in maps

We next evaluated the ability of Emap2lig-Find to detect ligand densities that are not modeled in the associated PDB entries. We examined two scenarios. In the first scenario, we selected EMDB map entries with associated PDB entries with multiple versions of structure files (as recorded in the PDB version archive mmCIF logs), in which the final version includes ligands absent or with shifted coordinates from the initial version. These differences reflect re-interpretation of the EM map and subsequent incorporation of additional ligands during model refinement. We applied Emap2lig-Find to the corresponding EM maps and assessed whether these newly added or shifted ligands could be detected. These ligands were not identified in the initial modeling, suggesting that they may be difficult to detect by visual inspection alone.

**Fig. 4a to 4c** are such maps with multiple versions of PDB files, and they are not included in our dataset. **Fig. 4a** is ligand detection results for a 2.33 Å resolution map of prostaglandin D2 receptor (EMD-38762). In addition to a phosphatidylinositol (A1D5Q) modeled by the authors in the initial version of the PDB file, Emap2lig-Find also identified the density of indomethacin (IMN) at the bottom of the map. Notably, this ligand was subsequently modeled by the authors in a later version of the PDB entry. The next map (**Fig. 4b**) is of oxoglutarate receptor 1 at resolution 2.84 Å embedded in the membrane system. The authors of this map initially modelled a long-chain fatty acid (A1D7P), that is seen on the left of the map. In addition, Emap2lig-Find detected long density, which turned out to be cholesterol in membrane (CLR) as modelled in the updated PDB file. A more complex case is illustrated in **Fig. 4c**, which shows a map of murine perforin-2 determined at 4 Å resolution (EMD-15086). In the initial deposition, both cyclohexyl-hexyl-β-D-maltoside (MA4) and N-acetylglucosamine (NAG) were modeled. The ligands in the outer circle were 16 MA4 molecules while the inner circle of ligands were 16 NAG molecules. In a subsequent revision, the authors removed MA4 entirely and repositioned the NAG coordinates. Emap2lig-Find detected no density supporting MA4 and instead identified density consistent with the revised NAG position, in agreement with the final deposited model.

**Fig. 4.**
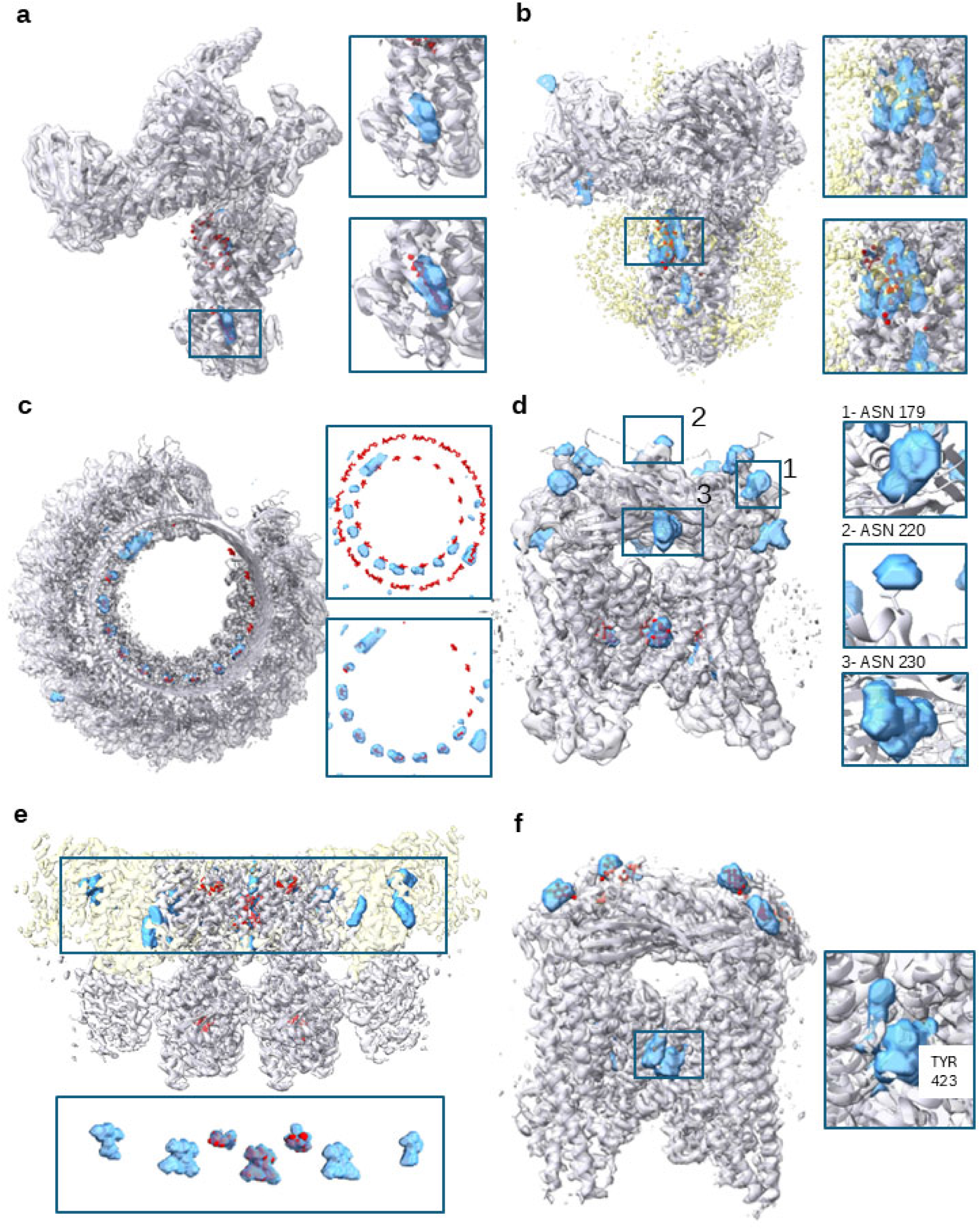
Detected ligand density in EM maps that are not modelled in PDB entries. Panels a–c show examples of detected ligands in EM maps associated with PDB entries that have two versions of deposited structural models. Ligands were identified in the maps but were absent in the initial PDB version, and they are consistent with ligands present in the later, updated version. The cryo-EM map is shown as gray density, and the deposited PDB structure is represented as a gray cartoon. The prediction by Emap2lig-Find in blue density and the deposited PDB structure of the ligand in red. Insets at the top show the initial deposited structures without the ligand, while those at the bottom show the updated structures with the modeled ligand from the later PDB version. **a**, prostaglandin D2 Receptor (PDB: 8XXV; EMD-38762; res.: 2.33 Å). Detected ligands: A1D5Q and IMN (indomethacin). Map recall using the second version of the PDB entry: 0.42. **b**, oxoglutarate receptor 1 (OXGR1) bound to leukotriene E4 and Gq proteins (PDB: 8YYX; EMD-39682; res.: 2.84 Å). Detected ligand: A1D7P and CLR (cholestrol). Map recall: 0.73. **c**, murin perforin-2 pore in twisted form (PDB: 8A1S; EMD-15086; res: 4 Å). Detected ligand: NAG. Map recall: 0.50. Panel d-f show the unmodeled ligands in the PDB entries. **d**, human TRPML1 channel structure in agonist-bound open conformation (PDB: 5WJ9; EMD-8841; res: 3.49 Å). Insets show predicted densities near the ASN residues 179, 220, and 230 of chain A. Emap2lig-Find identified 3 potentially NAG ligands in each of 4 chains of the protein alongside 1 copy of AQV ligand in each chain which was correctly annotated. **e**, STING oligomer (PDB: 8FLM; EMD-29282; res: 2.9 Å). A dimer is modelled in the PDB file. The cytosol is located on the lower side of the membrane, where cGAMP (CCD: 1SY) is binding, while the region above the membrane corresponds to the extracellular region. Emap2lig-find identified at least two more complete densities of Y6H ligands and 2 partial densities. **f**, TRPML3/ML-SA1 complex (PDB: 6AYF; EMD-7019; res: 3.62 Å). In addition to 4 NAG modelled in the PDB file, the method detected additional ligand densities near TYR423, which are most probably the ML-SA1.

The second set of examples concerns the detection of ligands that were not modeled by the original authors. We identified 17 structures in the test set that appear to contain unmodeled glycosylation sites, including six maps corresponding to transient receptor potential mucolipin 1 (TRPML1) channel proteins. In these cases, Emap2lig-Find predicted the presence of ligands located near the consensus sequence for N-linked glycosylation, Asn–X–Ser/Thr (X can be any amino acid except proline) in the upper part of the density in the figure. Based on this sequence context, the predicted ligand type is expected to be a carbohydrate moiety, such as N-acetylglucosamine (NAG). One such example is shown in **Fig. 4d** (EMD-8841; resolution 3.49 Å, PDB ID: 5WJ9). We observed twelve similar elongated densities near the consensus sequence at Asn179, Asn220, and Asn230 in all four chains of TRPML1, which were identified as putative ligand density. A literature survey revealed that Asn179, Asn220, and Asn230 are glycosylation sites that are important for the function of TRPML1 channels^6^. Therefore, we conclude that these densities are highly likely to correspond to carbohydrate moieties.

**Fig. 4e** shows a map of the STING dimer embedded in the membrane (EMD-29282; resolution: 2.9 Å; PDB: 8FLM). Although only a single dimer is modeled in the PDB structure, neighboring dimers are present on both sides, albeit with lower density. In the cytosolic domain, the complex binds two cGAMP molecules (CCD code: 1SY), which were correctly identified by Emap2lig-Find (ligand recall: 0.75 and 0.79). Within the membrane region, two copies of C53 ligand (CCD code: 9IM; the smaller compound shown in the inset panel) are identified, which is known to assist the higher-order oligomerization in the presence of the cGAMP and Y6H ligands, as well as one molecule of NVS-STG2 (CCD code: Y6H), a molecular glue that induces higher-order oligomerization between the dimers of the STING dimers^7^. In addition to the compounds modeled in the PDB structure, Emap2lig-Find detected four additional copies of Y6H at the interfaces between STING protein pairs, which is structurally plausible.

The last example is TRMPL3 depicted in **Fig. 4f** (PDB ID: 6AYF; EMD-7019; resolution 3.62 Å), which was labeled as the ML-SA1 (CCD code: AQV) compound bound structure^8^. The PDB entry contains 8 N-acetylglucosamine (NAG) molecules, corresponding to glycosylation moieties located at the top region of the structure, all of which were correctly detected. Emap2lig identified four additional densities near Tyr423 in all four protein chains, with similar shapes and consistent positions relative to the residue. Although the authors could not determine the orientation of ML-SA1 due to resolution limitations, they identified Tyr423 as a key binding residue. These densities therefore likely correspond to ML-SA1 molecules present in all four chains. Collectively, these examples demonstrate that Emap2lig-Find serves as an independent, density-driven validator of ligand annotations during structure modeling and would be highly useful for researchers.

### Building compound structures with Emap2ig-Build

We next evaluated Emap2lig-Build, the second stage of the Emap2lig pipeline, which reconstructs atomic models of compounds with known chemical identities from detected ligand densities. Evaluation was performed on cases in which Emap2lig-Find achieved an instance-level recall above 0.3, corresponding to 74.6% of the test cases (n = 1,040), as accurate ligand modeling is not practical when density detection is extremely limited. For a predicted ligand density, Emap2lig generates 64 conformations. **Fig. 5a** illustrates ligand conformations generated for a predicted density corresponding to ligand A1L04 (PDB 8Z3S; EMD-39753). Ten diverse conformations selected from 64 generated candidates are shown. The most accurate pose (thick yellow tube) achieved an atomic structural deviation of 0.77 Å relative to the deposited structure. The remaining conformations correctly positioned the three ring systems despite deviations of up to 5.5 Å.

**Fig. 5.**
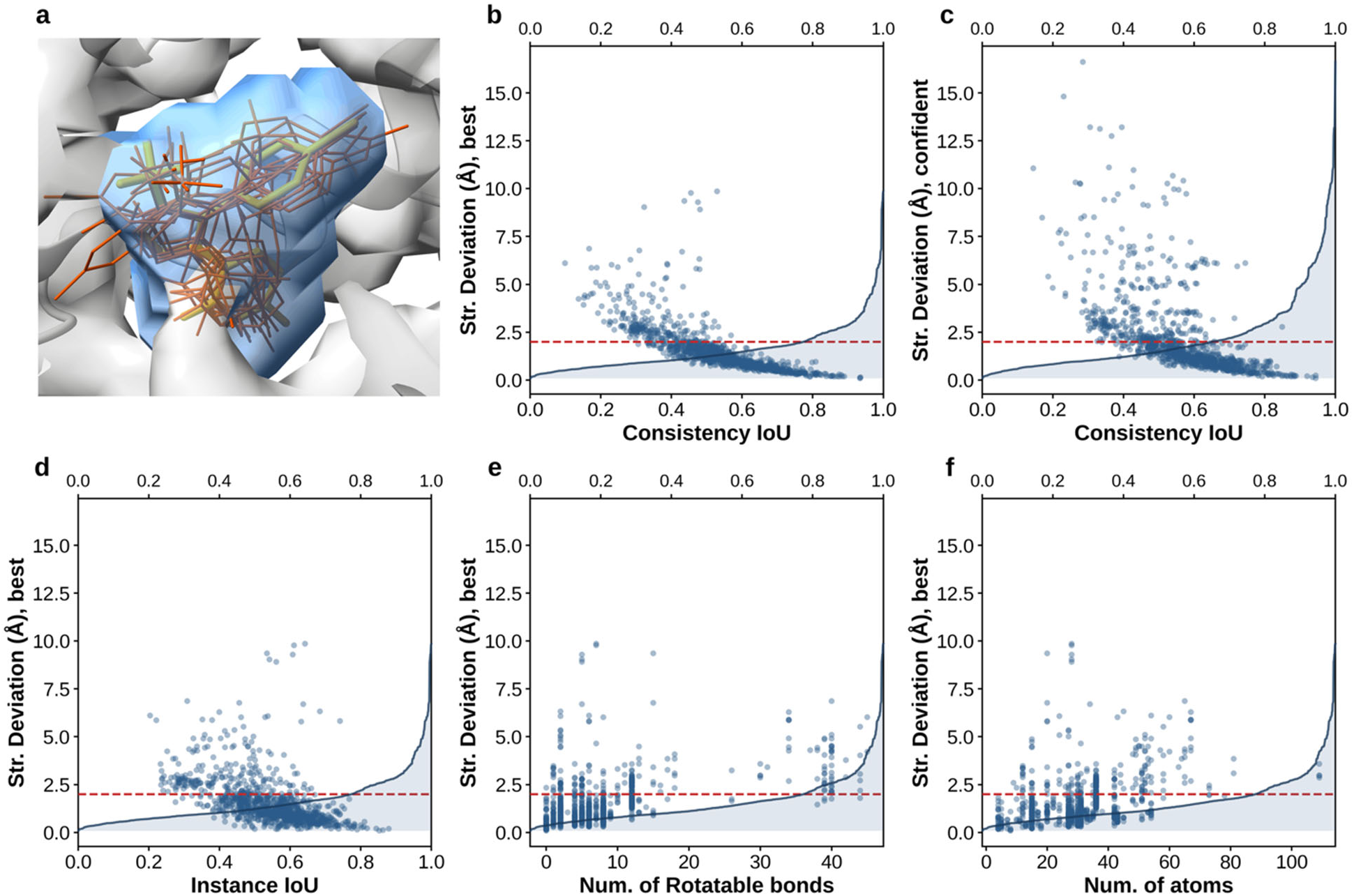
Evaluation of Emap2lig-Build. Structure modeling on the test-set of ligand instances with instance-level detection recall above 0.3 by Emap2lig-Find (n = 1040 instances). Red dashed horizontal lines in **b–f** mark y = 2.0 Å as a reference structure deviation level. Panel **b-f** overlays scatter points with the cumulative distribution of the vertical-axis quantity (structure deviation) on a secondary horizontal axis at the top of the panel. **a.** Representative structure modeling for ligand A1L04 (PDB ID: 8Z3S; EMD-39753). Ten sampled conformers spanning a structure deviation from 0.77 Å (best) to 5.5 Å (worst), shown inside the blue volume corresponding to the predicted instance density by Emap2lig-Build. The yellow conformer is the highest-quality sample (structure deviation: 0.77 Å to the reference structure). **b.** Consistency IoU versus structure deviation (Å) for the best conformer among 64 generated conformers (median structure deviation: 1.23 Å, mean: 1.57 Å; median consistency IoU: 0.53, mean: 0.53). **c.** The same metrics for the conformer with the highest consistency IoU among 64 sampled candidates (median structure deviation: 1.47 Å, mean: 2.17 Å; median consistency IoU: 0.57, mean: 0.56). **d.** Instance segmentation IoU from Emap2lig-Build versus structure deviation (Å) for the best conformer (median instance IoU: 0.55, mean: 0.54). Instance IoU measures overlap between the predicted and reference instance masks. **e.** The structure deviation (Å) for the best conformer versus the number of rotatable bonds inferred from RCSB chemical-component SMILES with RDKit (median: 6, mean: 8.6 for this subset; oligomeric ligands are summed across monomer components). **f.** Structure deviation (Å) for the best conformer versus the number of heavy atoms in the reference ligand (median: 31, mean: 30.7).

We start by evaluating modeling results in two key performance levels: In **Fig. 5b**, we evaluated the best (the model with the lowest structure deviation to the structure in the PDB file; or native) structure model among 64 generated models. In **Fig. 5c**, generated ligand conformations were sorted by a model confidence score, Consistency IoU, which we explain below. Individual data used in **Fig. 5** is provided in **Supplementary Table S8**.

The confidence of a generated ligand conformation was evaluated using the overlap between the modeled structure and the predicted ligand density, termed the Consistency Intersection over Union (IoU). The Consistency IoU measures the overlap between (i) a 3D mask of the modeled structure generated by placing 1.4 Å-radius spheres at atomic positions and (ii) the ligand density predicted by Emap2lig-Find. In **Fig. 5b**, the best model among 64 generated conformations and its Consistency IoU are shown for each target density. Overall, 77.3% of targets had at least one accurate model with a structural deviation below 2.0 Å (red dashed line). Consistency IoU correlated well with structural accuracy, particularly for models below 2.0 Å deviation. In a more practical setting, selecting the model with the highest Consistency IoU yielded a success rate of 65.0% (**Fig. 5c**). Considering multiple top-ranked conformations improved performance to 72.1%, 73.9%, and 75.8% within the top 5, 10, and 20 predictions, respectively. We identified a primary failure mode in 36 cases where the selected pose showed high confidence (Consistency IoU > 0.5) despite being structurally incorrect (deviation > 5.0 Å). In most of these cases, a flipped ligand orientation achieved a higher Consistency IoU than the correct pose, despite deviating more substantially from the native structure.

The quality of the initial segmentation influences downstream ligand modeling performance. In **Fig. 5d**, we examined the relationship between ligand modeling accuracy and ligand density detection accuracy, quantified by Instance IoU. Instance IoU measures the overlap between (i) the ligand density detected by Emap2lig-Find and (ii) the native ligand voxels derived from the deposited structure. As expected, modeling accuracy correlate with segmentation quality: 85.1% of ligands were reconstructed within 2.0 Å when the Instance IoU exceeded 0.4, whereas only 16.4% were modeled accurately when the Instance IoU was below 0.4.

In **Fig. 5e** and **5f**, we evaluated the effect of ligand complexity quantified by the number of rotatable bonds and atoms on modeling accuracy. The largest rotatable-bond count among successful predictions is 44, observed for a single case (EV9, a long-chain amine-modified phospholipid; PDB ID: 7DUW). For ligands with fewer than 15 rotatable bonds, Emap2lig-Build achieved a success rate of 83.2% within 2.0 Å (**Fig. 5e**). In contrast, performance decreases markedly with increasing flexibility, reaching 8.1% for ligands with more than 30 rotatable bonds. A similar trend is observed with ligand size: success reaches 82.9% for ligands with ≤40 atoms, whereas larger ligands show increased deviation and reduced accuracy (**Fig. 5f**).

Comparison of Emap2lig-Build with four existing methods, fitmap in ChimeraX^21^, VESPER^22^ in two scoring modes, the vector-product (VESPER’s original scoring scheme) and cross-correlation, and Phenix LigandFit^23,24^, which can model ligand conformations for a given ligand density (**Supplementary Table S9**), showed that Emap2lig-Build consistently outperformed these existing methods. As fitmap and VESPER perform ligand fitting, we generated ten conformers per ligand using OpenEye OMEGA^25^ and docked them. LigandFit from the Phenix package performs flexible ligand modeling. We generated ten conformers with LigandFit. Across three selection strategies (best conformer, most confident conformer by consistency IoU, and top-10 confident conformers), Emap2lig-Build consistently outperformed existing methods. Using the best generated conformer, it achieved a 77.3% success rate (mean deviation 1.57 Å; median 1.23 Å), compared with 65.0% (2.17 Å mean) when using only the most confident prediction. When top-10 confident conformations were considered, success elevated to 73.9% (1.74 Å mean), substantially exceeding competing methods, which showed mean deviations of 4.8–5.5 Å. Per-ligand analysis (**Supplementary Fig. 6, Supplementary Table S10**) further confirmed this trend, with Emap2lig-Build yielding lower deviations in >93% of cases across all baselines.

### Examples of modeled ligand conformations by Emap2lig-Build

**Fig. 6** shows representative examples of Emap2lig-Build ligand modeling. For each case, we display the cryo-EM map with Emap2lig-Find ligand density (blue), the lowest-deviation pose, the most confident pose, and the distribution of structural deviations across 64 generated conformers.

**Figure 6:**
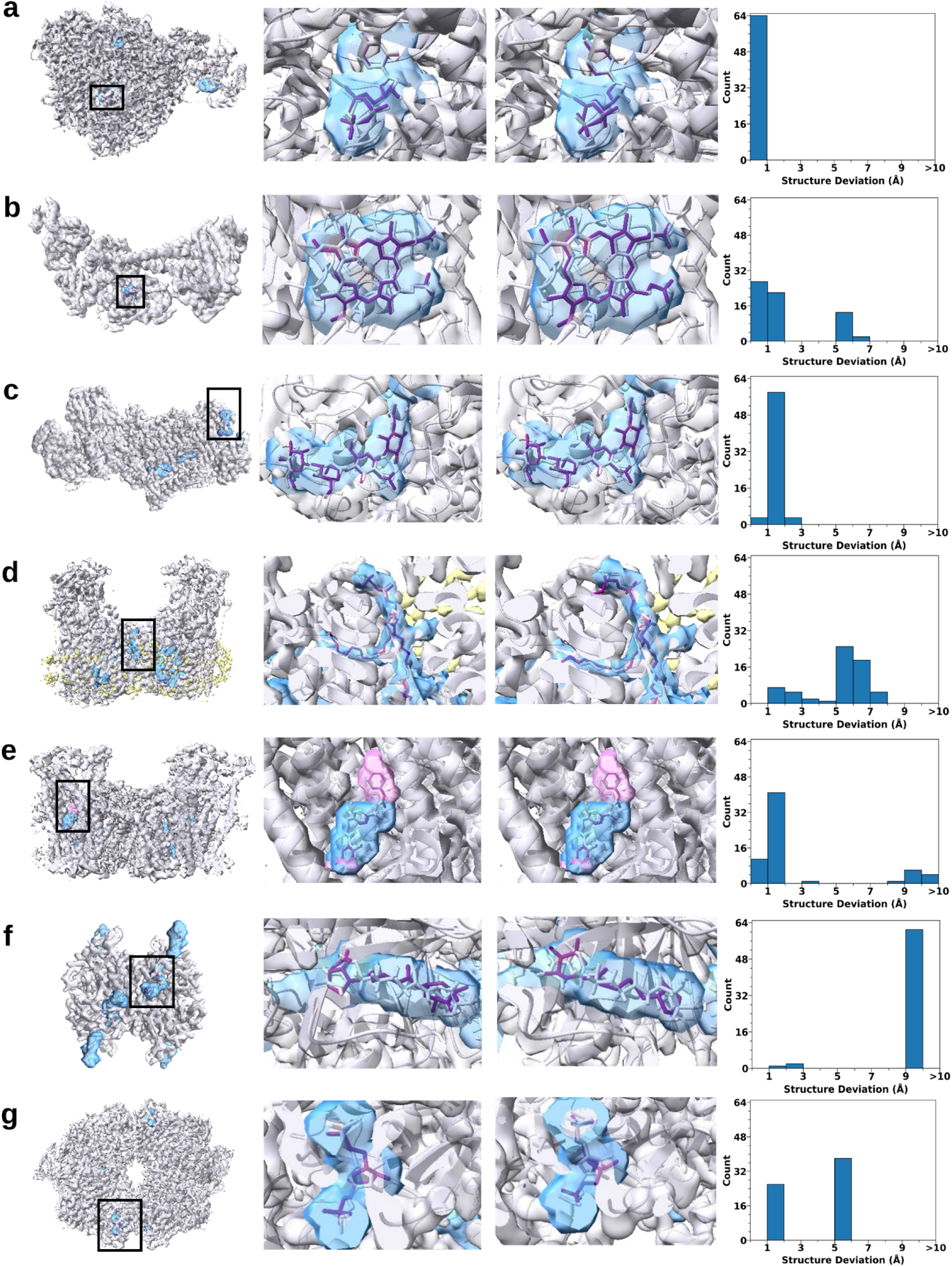
Examples of ligand modeling by Emap2lig-Build in cryo-EM density maps. Each row presents four panels: (left) the full cryo-EM map shown in gray density with all Emap2lig-Find predicted blobs highlighted in blue. The inset region marked by a black box shows the modeled ligand on interest shown on right columns. (center-left) close-up of the best conformation with the smallest structure deviation among 64 generated; (center-right) close-up of the most confident pose among 64 generated; (right) histogram structure deviation (Å) values of all 64 conformers modeled by Emap2lig-Build. Purple sticks show the modeled ligand pose; gray sticks show the native ligand from the deposited PDB structure. **a.** bat influenza A polymerase elongation complex (PDB: 6SZV; EMD-10355; res.: 2.5 Å). Ligand: 2KH; ligand recall: 0.96; the smallest str. deviation: 0.48 Å; str. deviation of the most confident conformation: 0.69 Å. **b.** guanylate cyclase (PDB: 6JT0; EMD-9883; res.: 4 Å)a. Ligand: HEM; ligand recall: 0.899; the smallest str. deviation: 0.63 Å; the most confident: 0.80 Å. **c.** Activated Drs2p-Cdc50p (PDB : 6ROJ; EMD-4974; res.: 2.9 Å). Ligand: NAG-NAG-BMA-BMA; ligand recall: 0.94; the smallest str. deviation: 0.87 Å; the most confident: 0.87 Å. **d.** multiple peptide resistance factor (PDB: 6LVF; EMD-0992; res.: 3.7 Å). Ligand: LHG; ligand recall: 0.94; the smallest str. deviation: 1.46 Å; the most confident: 1.53 Å. Yellow density represents the membrane region**. e.** pancreatic ATP-sensitive potassium channel (PDB: 5YKE; EMD-6831; res.: 4.11 Å). Ligand: GBM; ligand recall: 0.58; the smallest str. deviation: 0.54 Å; the most confident: 0.54 Å. The pink density represents the volume of the modelled ligand conformation. **f.** avidin with biotin-tempo (PDB: 7ZYL; EMD-15026; res.: 2.08 Å). Ligand: KFL; ligand recall: 0.66; the smallest str. deviation: 1.94 Å; the most confident: 9.72 Å. **g.** PKM2 protein (PDB: 6TTQ; EMD-10584; res.: 2.7 Å). Ligand: FBP; ligand recall: 0.98; the smallest str. deviation: 1.08 Å; the most confident: 5.85 Å.

The first example (**Fig. 6a**) shows a successful case on a high-resolution cryo-EM map of a bat influenza A polymerase elongation protein–RNA complex (EMD-10355, 2.5 Å). Emap2lig-Find predicted three ligand regions, correctly identifying two bound ligands and a third apparent false positive, which in fact corresponds to a disconnected guanine molecule, thus not entirely incorrect. The ligand of interest (2KH) was detected with high recall (0.96), enabling accurate modeling. Both the best and most confident poses achieved sub-angstrom accuracy (0.48 Å and 0.69 Å), closely matching the native structure, with minor deviations in phosphate-coordinating atoms. All 64 generated conformers were within 1 Å of the reference structure, indicating robust performance given an accurate initial detection.

Emap2lig-Build maintained strong performance at lower resolution. In a 4 Å map of a guanylate cyclase protein (EMD-9883) (**Fig. 6b**), Emap2lig-Find identified a single heme ligand (HEM) with high recall (0.899) and no false positives, enabling accurate modeling with best- and most-confident-pose deviations of 0.63 Å and 0.80 Å, respectively. The four pyrrole rings were correctly placed, with only minor overall displacement; although the deviation distribution broadened (up to ∼7 Å), most conformers remained within 1 Å.

The method also generalized to flexible glycans (**Fig. 6c**). In the activated Drs2p–Cdc50p complex (EMD-4974, 2.9 Å), Emap2lig-Find detected a four-residue glycan chain (two NAG and two BMA) with recall of 0.94, despite several small false positives that can be readily distinguishable due to their small size. The best and most confident conformers were identical (0.87 Å deviation), and most of the 64 generated poses fell within 1–2 Å, indicating robust near-native sampling.

The next example **(Fig. 6d**) is a challenging membrane protein target, a multiple peptide resistance factor (EMD-0992, 3.7 Å). Of nine predicted ligand densities, six were initially classified as false positives; however, four displayed elongated shapes consistent with lipid acyl chains, suggesting unmodeled lipids rather than true false detections, while the remaining two may correspond to small lipid fragments of uncertain identity. The target ligand (LHG), a phospholipid, was detected with high recall (0.94). Although conformational flexibility of the lipid tails led to deviations >5 Å for most conformers, Emap2lig-Build still identified a highly accurate pose as the most confident prediction (1.53 Å) and produced an even closer alternative within the conformer pool (1.46 Å).

**Fig. 6e** illustrates a case where Emap2lig-Build compensates for incomplete detection by Emap2lig-Find. In the pancreatic ATP-sensitive potassium channel (EMD-6831, 4.11 Å), 17 regions were identified, 13 of which were initially labeled as false positives; however, most exhibited elongated lipid-like densities consistent with unmodeled membrane lipids, while only four appeared to be true false positives. The target ligand (GBM), a hydrophobic inhibitor, was partially detected with recall of 0.58, capturing only a fraction of the ligand density (blue region). Despite this incomplete initialization, Emap2lig-Build expanded the detected region during modeling (pink) and recovered the full ligand, achieving a best-pose deviation of 0.54 Å, with identical performance for the most confident pose. Although a subset of conformers showed large deviations (>8 Å), the majority (53/64) clustered within 2 Å, indicating robust recovery from partial detections.

**Fig. 6f** shows a case where a near-native pose was generated but not selected as the most confident prediction. In avidin bound to biotin-tempo (EMD-15026, 2.08 Å), Emap2lig-Find predicted nine regions (one false positive) and achieved a ligand recall of 0.66 for KFL. Although a correct pose was present among the 64 conformers (best deviation: 1.94 Å), the most confident pose corresponded to a flipped orientation with a large deviation (9.72 Å), despite similar consistency IoU between the two orientations. The ensemble was dominated by the flipped state, with most conformers clustering above 9 Å.

**Fig. 6g** presents a similar failure case in pyruvate kinase 2 (EMD-10584, 2.7 Å). Emap2lig-Find predicted nine regions with three false positives and achieved high recall (0.98) for FBP. While the best pose was accurate (1.08 Å), the most confident pose showed substantial deviation (5.85 Å) due to misalignment of the central ring, which propagated through the molecule. The conformer distribution was bimodal, with clusters below 2 Å and at 5–6 Å, indicating coexistence of correct and systematically misaligned solutions.

## Discussion

Emap2lig is a two-stage deep learning framework for automated ligand density detection (Emap2lig-Find) and 3D atomic coordinate modeling (Emap2lig-Build) directly from experimental cryo-EM maps, without requiring prior ligand localization or a macromolecular model. This distinguishes it from existing approaches that typically depend on well-resolved protein structures and predefined binding pockets. By operating directly on experimental maps, Emap2lig is well suited for early-stage structure determination of protein–ligand complexes where binding sites may be unknown, transient, or only partially resolved. Its ability to identify and model ligands without structural priors further enables unbiased interpretation of cryo-EM density, facilitating discovery of cofactors, lipids, metabolites, post-translational modifications, and drug-binding events. This is particularly valuable for structural biology and drug discovery applications, where rapid elucidation of ligand-binding modes can accelerate mechanistic insight and structure-guided design.

A key challenge in developing Emap2lig is that ligands in deposited PDB structures are often incompletely annotated, particularly for membrane-associated lipids and glycans such as NAG that occur in multiple copies in cryo-EM maps. Consequently, distinguishing false positives from genuine but unmodeled ligand densities is not always straightforward. This limitation also highlights a potential application of Emap2lig in assisting structural biologists to identify and annotate previously unmodeled ligands. This capability is particularly relevant for membrane proteins, where lipid environments are frequently absent or incomplete in PDB models, limiting downstream computational modeling and simulation.

Another area for improvement is pose ranking in Emap2lig-Build. Our analyses indicate that near-native conformations are frequently present within the generated ensemble, but selecting the optimal pose remains challenging in some cases. Pose ranking currently relies primarily on geometric overlap with the predicted ligand density; incorporating learned scoring functions using deep learning may improve confident pose selection and reduce orientation-related failure modes. A further direction is integration with macromolecular structure prediction, as recent advances in protein and nucleic acid modeling^26–28^ increasingly enable unified frameworks for biomolecular structure determination directly from cryo-EM maps.

Overall, these results highlight Emap2lig as not only a ligand discovery and modeling platform, but also a potential quality-control tool for deposited cryo-EM structures. We anticipate that it will facilitate ligand annotation and accelerate structural interpretation of cryo-EM data.

## Author Contributions

D.K. conceived the study. S.L. contributed to code implementation. S.L. conducted model training. A.J., S.L., Y.K contributed to dataset preparation. S.L., A.J, D.K. analyzed the data. S.L. and A.J. drafted the paper. A.J. analyzed individual cases. S.L. implemented the web GUI. J.H.P. implemented it on the web server. D.K. critically edited it.

## Acknowledgements

This work was partly supported by the National Institutes of Health (R35GM158267, R21AI187928) and the National Science Foundation (IIS2211598, DMS2151678, DBI2146026 and DBI2422620).

## Methods

### MUNet for processing an input cryo-EM map

To process volumetric cryo-EM density maps and extract structural features at multiple spatial scales, the Emap2lig pipeline employs two instances of an enhanced U-Net architecture^14^ (**Fig. 1a**, blue and orange boxes), referred to as the Modern U-Net (MUNet). The MUNet is derived from the original 3D U-Net framework^14^ and is adapted to model both local density patterns and long-range structural context within macromolecular maps. The architecture follows a standard encoder–decoder structure with skip connections (**Supplementary Fig. 1**). In the upper levels of the network (Convolutional Encoder Blocks 1–2 and Convolutional Decoder Blocks 1–2), feature extraction relies exclusively on 3D convolutional layers. These layers capture localized density variations, including edges, gradients, and compact structural motifs, which are spatially constrained and benefit from translation-equivariant operations. On the other hand, the lowest-resolution stages of the network (Attention Encoder Block 3, Middle Block, and Attention Decoder Block 3) incorporate full self-attention blocks^29^. At this compressed feature scale, attention enables global receptive fields and facilitates modeling of long-range structural relationships within the macromolecular assembly. Restricting attention to low-resolution feature maps preserves global context while avoiding the quadratic computational cost that would arise from applying attention directly to high-resolution volumetric inputs.

We further modernized the backbone architecture to improve optimization stability. ReLU activations were replaced with the SiLU^30^ function to enable smoother gradient propagation. In addition, 3D Batch Normalization was replaced with Group Normalization^31^ (with a group size of 32). Because volumetric density tensors impose strict memory constraints and limit batch sizes during training, Group Normalization provides more stable behavior under small-batch regimes.

The two MUNet instances are configured with distinct spatial and channel capacities to match their functional roles. The first MUNet (**Fig. 1a**, blue box) processes 64 Å³ input patches with 32 base feature channels and is optimized for broader contextual segmentation. The second MUNet (**Fig. 1a**, orange box) operates on 48 Å³ patches with 64 base channels, increasing representational capacity to resolve finer-grained semantic chemical labels. Additional architectural details are provided in **Supplementary Information 1**.

### Conformer embedder and pairformer for single- and pair- representations

The conformer embedder in Emap2lig-Build translates a reference ligand conformer into a structured representation composed of single-atom and pairwise features. It ingests three inputs: per-atom features describing elemental and chemical properties, pairwise features encoding atom–atom relationships, and the 3D coordinates of the reference geometry. Binary atom and pair masks are provided to distinguish valid atoms from padding introduced for batching. **Supplementary Information 2** provides more details.

The conformer embedder first projects the input atom features into a latent space using a linear layer (**Supplementary Information 3)**. The atom features are linearly converted with valid atom mask to produce the single atom representation. Computation of pair features starts from the initial atom representation and augments it with geometric information computed from the reference coordinates. Specifically, it derives relative position vectors and inverse-square interatomic distances for each atom pair. These geometric terms, together with a learned embedding of the pair mask, are linearly projected and added to the pair representation. Finally, the updated atom embeddings are projected into the pair feature dimension and added to the corresponding pair representations (for both atoms in each pair) (initial pairwise representation).

The pairformer refines initial single and pairwise representations from conformer embedder and serves as the central module for chemical feature propagation (**Supplementary Information 4**). It follows the architectural design of AlphaFold3^32^, with the number of blocks reduced to four. Each block begins with a pair-to-atom attention layer, in which pairwise representations guide the update of single-atom representations. An atom-to-pair outer product layer then projects the updated single-atom representations back into pair space, allowing pairwise features to reflect the evolving atomic states. Triangle multiplication and triangle attention layers operate over triplets of atoms to enforce geometric consistency across three-body relationships. Finally, transition layers consolidate information within both single-atom and pairwise representations. Binary masks are applied at each stage to maintain valid structural support and remove contributions from padding.

### Instance Segmentation Module

The instance segmentation module predicts a ligand-specific 3D probability mask to isolate the target ligand instance within the cryo-EM map (**Supplementary Information 5, Supplementary Fig. 2**). The module receives three inputs that correspond to the left-side inputs in **Supplementary Fig. 2**: (i) the voxel feature map *F* (blue box), extracted by the second MUNet (**Fig. 1a**, orange box); (ii) the atom features, the single-atom representation *s_i_* with mask 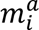 from the pairformer; and (iii) the prompt point *p*, a 3-D coordinate sampled from a ligand region detected by Emap2Lig-Find. To distinguish the target ligand from adjacent proteins, nucleic acids, or other ligand instances, the module uses a prompt point as a spatial anchor. This point is stochastically sampled from the ligand density identified in the Emap2lig-Find stage. Conditioned on this localized coordinate prior and the reference chemical representations, the module refines a ligand-specific mask within the full map context, yielding an instance-level segmentation of the target molecule. Based on the refined mask, 8,192 points are sampled from the masked volume to construct point-wise features for downstream structure generation, while global features are extracted via adaptive 3D pooling to capture the overall spatial context of the ligand region. The prompt point is converted to per-voxel relative-position map, which conditions segmentation on a specific spatial location. These inputs are processed through four sequential AtomsContextBlocks (the four stacked blocks inside the dotted rectangle), which progressively refine the volumetric representation.

At each AtomsContextBlock, the relative position map *r*_*rel*_, and the atom features (*s_i_* with mask 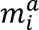) are fed in alongside the voxel feature map, matching the three incoming arrows in Supplementary Fig. 2. The voxel feature map is first processed by local 3-D convolutions and downsampled by a factor of four to reduce spatial resolution and enable cross-attention at lower computational cost. The downsampled voxel features, augmented with the encoded *r*_*rel*_, serve as queries, while the atom features (*s_i_*, 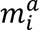) serve as keys and values. This design allows each spatial location to attend to atom-level context, thereby conditioning the segmentation on ligand-specific chemical information guided by the prompt point.

After cross-attention, the refined latent features are upsampled back to the original spatial resolution. In parallel, a residual pathway (the dashed arrow in **Supplementary Fig. 2**) composed of local 3-D convolutional layers bypasses the attention branch and preserves high-frequency spatial information. The upsampled attention features and residual convolutional features are combined to form the updated voxel representation. After four such refinement stages, a final 3D convolution (the Segmentation Head in **Supplementary Fig. 2**) projects the features to a single-channel output, producing the voxel-wise instance probability mask *seg* (*P*) (dashed box with a generated mask in pink) that delineates the target ligand instance.

Following mask prediction, the backbone voxel features (blue box on the left), the augmented detection predictions (red box on the left; *y_aug_*, carried over from the segmentation head of Emap2Lig-Build U-Net), and the predicted instance mask (*seg*, *P*) are concatenated and passed through the ConvBlock to produce the fused voxel feature *V* (orang box at the right bottom). This fusion integrates backbone representations, detection cues, and instance-level spatial focus into a unified feature space for conditioning the diffusion module. Based on the refined mask, TopK selects the *K* = 8,192 highest-probability voxels to construct the point output used for structure generation.

In addition, a global feature vector is extracted from the fused voxel feature *V*. Specifically, world coordinate information (the per-voxel *xyz* coordinates computed from the map origin *o* and voxel size δ, as part of the Voxel Feature in Supplementary Fig. 2) is concatenated with *V* and transformed through a 1 × 1 × 1 convolution. The resulting features are modulated by a learned transformation of the predicted mask *P*, and 3-D adaptive average pooling is applied over the masked volume to obtain a permutation-invariant global feature vector *g* summarizing the segmented density distribution. In parallel, The top-K sampling on the instance probability mask with the *K* = 8,192 highest-probability voxels’ fused-feature vectors and their world coordinates are combined to form the point output (*v_k_*, *c_k_*). Both the point features (*v_k_*, *c_k_*) and the global vector *g* are then passed to the diffusion module (**Supplementary Fig. 3**; Algorithm 10) as ligand-specific spatial conditioning.

### Diffusion Module

The diffusion module (**Supplementary Information 6, Supplementary Fig. 3**) generates the final 3D atomic coordinates of the ligand using an Elucidated Diffusion Model (EDM) ^33^. Starting from Gaussian noise in coordinate space, the model iteratively denoises the atomic positions to produce a physically consistent molecular geometry. The denoising trajectory is conditioned on the single-atom and pairwise representations from the pairformer, as well as the predicted instance mask and voxel features from the instance segmentation module. This design couples internal chemical representations with external density evidence throughout the generative process.

The network consists of eight identical conditioning blocks. Within each block, single-atom representations are updated through self-attention with pair bias, where the pairwise representations modulate attention scores to enforce chemically consistent interactions. To incorporate spatial information from the cryo-EM map, single-atom representations attend to point-wise features at the sampled spatial locations through cross-attention, allowing each atom to integrate local density evidence. The global feature vector extracted from the instance segmentation module is broadcast to all atoms through a transition layer, enhancing their awareness of the overall density distribution.

A final coordinate decoder predicts denoising updates to the atomic positions at each diffusion step. Repeated application of these conditioned updates yields the final 3D molecular structure.

### Dataset Construction

The dataset was constructed from all entries in EMDB with associated PDB entries containing ligands, as of August 1, 2024. Structures with resolutions up to 6 Å were selected, as we anticipated developing a model capable of recognizing ligands up to this resolution. Ligands were defined as small molecules distinct from protein or nucleic acid chains, as well as saccharides covalently attached to protein side chains (e.g., NAG, BMA, and MAN), each containing more than three heavy atoms.

To assess the quality of fit between the modeled biomolecular complex and its ligands, the cross-correlation (CC) between the structures and the cryo-EM density map was calculated. For this, the density of the protein–ligand complex was first segmented using a 4 Å zoning radius, and ligand-only densities were segmented with a 2 Å radius. The smaller radius for ligands was chosen to minimize the inclusion of adjacent protein or nucleic acid density. Preliminary analysis indicated that ligand density was better defined and produced higher CC values at 0.5 × the author-recommended contour level. Consequently, this 0.5 × level was adopted for all ligand-only CC calculations, while the full complex used the 1.0 × author-recommended level.

The dataset was then constructed by applying two sequential filters. First, entries were retained if they exhibited a CC greater than 0.65 for both the full biomolecular complex and the ligand-only regions. Second, to ensure structural completeness, any entry containing one or more ligands with an atom-resolved fraction below 0.8 was excluded. The final filtered dataset comprises 7,471 EM–structure pairs, containing 94,379 ligand instances.

To create non-redundant data splits, protein and nucleic acid sequences were clustered. Protein chains were clustered using the Needleman-Wunsch alignment algorithm^34^ with a 25% sequence identity threshold. PDB structures were then grouped if any of their chains resided in the same cluster. A similar process was applied to nucleic acid sequences using an 80% sequence identity threshold. This resulted in a training set of 7,065 EMDB-PDB pairs (from 332 non-redundant clusters) with 91,814 ligand instances; a test set of 228 pairs (38 clusters) with 1394 instances; and a validation set of 178 pairs (41 clusters) with 1171 instances.

For each experimental map, the voxel spacing was first unified to 1.0 Å by applying trilinear interpolation of the electron density. The maps were then normalized by first replacing negative density values with 0.0 and setting half of the author-recommended contour level as the threshold for minimum density values. The density values were then normalized to the range of [0.0, 1.0] using the 98th percentile of non-zero values as the upper bound to handle outliers. We generated separate simulated density maps for each structural label. The structural labels include protein backbone and side chain atoms, phosphate, sugar, and base components of nucleic acids, as well as ligands for Emap2lig-Find. For Emap2lig-Build, the labels further include these categories along with six elemental types (C, N, O, P, S, and metals) and ring systems containing four, five, or six members. We assigned each voxel to a structural class based on the presence of heavy atoms within 1.414 Å of the voxel center. This distance threshold was chosen empirically to preserve fine structural features, such as the hollow regions in ring systems. Simulated density maps were generated using the TEMPy2^35^ algorithm for each structural classes.

The training dataset of Emap2lig-Build was constructed based on the outputs of the Emap2lig-Find stage. Specifically, ligand instances were retained only if they achieved an instance-level recall ≥ 0.3 during Emap2lig-Find. This filtering resulted in 72,938 ligand instances out of 94,379 total instances (∼77.3%), spanning 1,293 distinct ligand classes out of 1,652 total classes. By restricting the build-stage training data to ligands that are at least partially localized in the find stage, this strategy reduces noise from the dataset. For the Emap2lig-Build stage, we adopted the same training/validation/testing split at the map level as used in Emap2lig-Find. All eligible ligand instances from each map were included in the corresponding split to construct the Emap2lig-Build dataset, ensuring no data leakage across splits.

The dataset included 1,807 ligand types. The number of heavy atoms of the ligands in our dataset ranged from 4 (with ligands such as CO3 (Carbonate)) to 200 (with ligands such as JSG) (average size: 32.4).

### Ligands with biological relevance

To define a list of biologically relevant ligands, we examined the literature associated with the PDB structures in which the ligands were present. Ligands were considered biologically relevant if the corresponding publications explicitly discussed their structural or functional roles, such as acting as inhibitors or cofactors. In addition, we consulted the BioLip2 database database^36^, which provides a curated definition of biologically irrelevant ligands. These ligands are primarily additives used during structure determination, such as buffer components, small ions, and lipids.

BioLip2 classifies buffer reagents, small ions, and lipids as biologically irrelevant. For ligands that were absent from the BioLip2 dataset we mainly perused literature to determine if they were relevant to the studies. There were 117 ligands shared between our irrelevant list and the BioLip2 irrelevant list. Some ligands listed as irrelevant in BioLip2 were considered relevant in our dataset because the associated publications discussed their biological roles; examples include TAU (taurine) and FOL (folic acid). Conversely, ligands classified as irrelevant in our dataset but absent from the BioLip2 irrelevant list included additional phospholipids (e.g., A1AF7) and saccharides such as MAN (α-D-mannopyranose).

In total, 1,343 ligands were labeled as biologically relevant, while 464 ligands were classified as irrelevant. The list of ligands is provided in Supplementary Tables S10 and S11.

### Maps with membrane systems

Among the maps in the test dataset, there were 109 maps for biological systems embedded in membrane. These maps are specified in the **Supplementary Table S2**. Maps with membrane were identified by manual inspection.

### Model Training

The overall training pipeline consists of two main stages: Emap2lig-Find and Emap2lig-Build. Emap2lig-Build further proceeds in two phases: pretraining and full (end-to-end) training.

### Emap2lig-Find Training Data Sampling

The training dataset was constructed from a curated list of EMDB/PDB entries. For each entry, we first generated a pool of 3D patches from the unified cryo-EM map. Patches were generated using a sliding window with a patch size of 80 Å³, and a stride of 40 Å in each dimension, as defined in the data generation configuration.

To enrich the dataset with relevant structural features and mitigate class imbalance, patches were saved selectively based on their content. A patch was included in the training pool if it satisfied one of two conditions: 1) If the patch contained at least 50% of the total voxels belonging to a single ligand instance (ligand-centric sampling). Or 2) If at least 10% of the patch’s total volume was occupied by voxels from any of the defined labels (backbone, sidechain, sugar, phosphate, base, or ligand) (General-object sampling). This process created an on-disk dataset of 301,843 voxel patches, from which the data loader sampled during training.

### Feature Augmentation

Feature augmentation was performed on-the-fly during the training process. From each 80 Å³ data patch loaded from disk, a smaller 64 Å³ subpatch was extracted from a random spatial location. This random cropping strategy served as a primary spatial augmentation, ensuring the model was exposed to diverse sub-regions of the input features. Furthermore, these cropped subpatches were subjected to additional geometric transformations, with a chance of being randomly mirrored along each of the three principal axes to further expand the diversity of the training data.

### Optimizer and Learning Rate Scheduler

We employed the ADOPT^37^ with an initial learning rate of 1 × 10^-4^, with default hyperparameters (betas = (0.9, 0.9999), ɛ =1 × 10^-6^). To manage the learning rate schedule, a ReduceLROnPlateau scheduler was utilized. This scheduler monitored the training loss, reducing the learning rate by a factor of 0.5 if no improvement was observed for 5 consecutive epochs.

The total loss function combines a segmentation loss and a regression loss with equal weights. The segmentation loss is the Dice Loss between output probability and target labels. The regression branch predicts the simulated density values for each channel corresponding to structural labels at every voxel. It is trained using the mean squared error (MSE) computed per channel between the predicted densities and the corresponding target simulated density maps. The regression outputs represent continuous density values (not probabilities) and are therefore not constrained to the range [0, 1]. The auxiliary heads used during training of the diffusion module are detailed in **Supplementary Information 7**. Further training details of Emap2lig-Find training are given in **Supplementary Information 8**.

The model was trained for 200 epochs with an effective batch size of 192 (batch size of 24 with 8× gradient accumulation) using the PyTorch Lightning framework. All experiments were conducted on a single NVIDIA A100 80GB GPU. To accelerate computation, TF32 matrix operations were enabled, and the model graph was further optimized through Torch compilation. Validation was performed at 5 epoch intervals throughout training.

### Emap2lig-Build Training

For more details, see **Supplementary Information 9**. The structure model training proceeded in two stages: pretraining and full (end-to-end) training.

In the pretraining stage, the conformer embedder, pairformer, instance segmentation module, and auxiliary module (**Supplementary Information 7**) were trained jointly. The training signals included the multi-class segmentation loss for the U-net, instance segmentation loss for instance segmentation module, as well as the auxiliary distogram loss and auxiliary classification loss from the auxiliary module. This stage establishes robust structural and density-aware representations before introducing the diffusion objective.

In the full (end-to-end) training stage, all components, including the diffusion module, were optimized jointly. The overall objective combines coordinate denoising losses with segmentation losses and auxiliary losses. The loss weights for both pre-training and full training are provided in **Supplementary Table S12 and S13**.

All training was conducted using the PyTorch Lightning framework with distributed data parallelism on eight NVIDIA H100 80 GB GPUs. Mixed-precision training (bf16) with TF32 support was enabled.

### Optimizer and Scheduler

For pretraining, we employed the ADOPT optimizer with a learning rate of 1 × 10^-3^, β values of (0.9, 0.95), and *ε* = 1 × 10^-8^. The learning rate was governed by a custom step-wise scheduler consisting of a linear warmup from 0 to the maximum learning rate over 2,000 steps, followed by a plateau, after which the learning rate decayed exponentially by a factor of 0.8 every 5,000 steps starting at step 7,000. Each GPU processed a batch of 2 ligand instances with gradient accumulation over 16 steps, yielding an effective batch size of 256 across 8 GPUs.

For full (end-to-end) training, the optimizer, learning rate schedule, and hardware configuration were the same as in the pretraining stage. The model was trained for up to 500 epochs with validation performed at 1-epoch intervals.

During inference, the model generates multiple candidate conformers per ligand, each conditioned on a different prompt point. To select the best conformer without access to native coordinates, we compute a consistency IoU score that measures the agreement between the predicted structure and the predicted instance segmentation mask. The candidate with the highest consistency IoU is selected as the final predicted conformer.

### Evaluation for Emap2lig-Build

During inference, the model generates *M* (=64) candidate conformers per ligand, each conditioned on a different prompt point. To select the most confident conformer without access to the correct coordinates, we compute a consistency IoU score that measures the agreement between the predicted structure and the predicted instance segmentation mask.

For each candidate conformer *m*:

1. The predicted atom coordinates 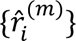 are converted into a binary voxel mask by labeling all voxels within a radius of 1.4 Å of any predicted atom position, producing a coordinate-derived mask 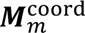.
2. The instance segmentation module independently predicts a density-derived probability mask ***P***^*(m)*^ for the same prompt point.
3. The consistency IoU is computed as the intersection-over-union between the coordinate-derived mask and the thresholded density-derived mask:

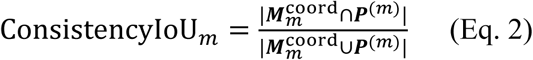

A high consistency IoU indicates that the predicted atom positions occupy the same spatial region as the predicted instance mask, meaning the coordinate prediction is geometrically consistent with the density evidence. The candidate with the highest consistency IoU is selected as the final predicted conformer.

Consistency IoU exploits the fact that two independent prediction pathways (coordinate generation via diffusion and spatial localization via instance segmentation) should agree when the model is confident.

### Structure deviation to the conformation in the PDB file

We evaluate the agreement between the predicted ligand conformation and the deposited PDB conformation without applying any rigid-body alignment (i.e., no rotation or translation). We refer to this alignment-free, map-frame metric as the *structure deviation*, to distinguish it from the conventional RMSD, which is usually computed after optimal superposition. Given a one-to-one correspondence between atoms in the predicted and reference structures, the structure deviation is defined as:

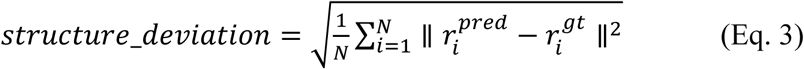

where 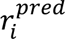 is the predicted coordinate of atom *i*, 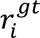 is the ground truth (correct) coordinate, and *N* is the total number of atoms, all evaluated in the cryo-EM map frame.

## Reference

1. Rubach, P. et al. Advances in cryo-electron microscopy (cryoEM) for structure-based drug discovery. Expert Opinion on Drug Discovery 20, 163–176 (2025).

2. Van Drie, J. H. & Tong, L. Cryo-EM as a powerful tool for drug discovery. Bioorganic & Medicinal Chemistry Letters 30, 127524 (2020).

3. Farheen, F., Terashi, G., Zhu, H. & Kihara, D. AI-based methods for biomolecular structure modeling for Cryo-EM. Current Opinion in Structural Biology 90, 102989 (2025).

4. Wagner, T. et al. SPHIRE-crYOLO is a fast and accurate fully automated particle picker for cryo-EM. Commun Biol 2, 218 (2019).

5. Bepler, T. et al. Positive-unlabeled convolutional neural networks for particle picking in cryo-electron micrographs. Nat Methods 16, 1153–1160 (2019).

6. Zhong, E. D., Bepler, T., Berger, B. & Davis, J. H. CryoDRGN: reconstruction of heterogeneous cryo-EM structures using neural networks. Nat Methods 18, 176–185 (2021).

7. Li, S., Terashi, G., Zhang, Z. & Kihara, D. Advancing structure modeling from cryo-EM maps with deep learning. Biochemical Society Transactions 53, 259–265 (2025).

8. Terashi, G., Wang, X., Prasad, D., Nakamura, T. & Kihara, D. DeepMainmast: integrated protocol of protein structure modeling for cryo-EM with deep learning and structure prediction. Nat Methods 21, 122–131 (2023).

9. Su, B., Huang, K., Peng, Z., Amunts, A. & Yang, J. CryoAtom improves model building for cryo-EM. Nat Struct Mol Biol 10.1038/s41594-025-01713-3(2025) doi:10.1038/s41594-025-01713-3.

10. Terashi, G., Wang, X., Maddhuri Venkata Subramaniya, S. R., Tesmer, J. J. G. & Kihara, D. Residue-wise local quality estimation for protein models from cryo-EM maps. Nat Methods 19, 1116–1125 (2022).

11. Robertson, M. J., Van Zundert, G. C. P., Borrelli, K. & Skiniotis, G. GemSpot: A Pipeline for Robust Modeling of Ligands into Cryo-EM Maps. Structure 28, 707–716.e3 (2020).

12. Sweeney, A., Mulvaney, T. & Topf, M. ChemEM: flexible docking of small molecules in Cryo-EM structures using difference maps. 2023.03.13.532279 Preprint at 10.1101/2023.03.13.532279 (2023).

13. Muenks, A., Zepeda, S., Zhou, G., Veesler, D. & DiMaio, F. Automatic and accurate ligand structure determination guided by cryo-electron microscopy maps. Nat Commun 14, 1164 (2023).

14. Falk, T. et al. U-Net: deep learning for cell counting, detection, and morphometry. Nat Methods 16, 67–70 (2019).

15. Greg Landrum et al. RDKit: Open-source cheminformatics. Zenodo 10.5281/ZENODO.591637 (2026).

16. Williams, D. C. & Inala, N. Physics-Informed Generative Model for Drug-like Molecule Conformers. J. Chem. Inf. Model. 64, 2988–3007 (2024).

17. The wwPDB Consortium. EMDB—the Electron Microscopy Data Bank. Nucleic Acids Res 52, D456–D465 (2024).

18. Bekker, G.-J. et al. Protein Data Bank Japan: Computational Resources for Analysis of Protein Structures. Journal of Molecular Biology 437, 169013 (2025).

19. Li, C. H. & Tam, P. K. S. An iterative algorithm for minimum cross entropy thresholding. Pattern Recognition Letters 19, 771–776 (1998).

20. Hattori, M., Tanaka, N., Kanehisa, M. & Goto, S. SIMCOMP/SUBCOMP: chemical structure search servers for network analyses. Nucleic Acids Res 38, W652–W656 (2010).

21. UCSF ChimeraX: Tools for structure building and analysis - Meng - 2023 - Protein Science - Wiley Online Library. https://onlinelibrary.wiley.com/doi/10.1002/pro.4792.

22. Han, X., Terashi, G., Christoffer, C., Chen, S. & Kihara, D. VESPER: global and local cryo-EM map alignment using local density vectors. Nat Commun 12, 2090 (2021).

23. Terwilliger, T. C., Klei, H., Adams, P. D., Moriarty, N. W. & Cohn, J. D. (IUCr) Automated ligand fitting by core-fragment fitting and extension into density. https://journals.iucr.org/d/issues/2006/08/00/dz5066/index.html.

24. Terwilliger, T. C., Adams, P. D., Moriarty, N. W. & Cohn, J. D. (IUCr) Ligand identification using electron-density map correlations. https://journals.iucr.org/d/issues/2007/01/00/ba5092/index.html.

25. Hawkins, P. C. D., Skillman, A. G., Warren, G. L., Ellingson, B. A. & Stahl, M. T. Conformer Generation with OMEGA: Algorithm and Validation Using High Quality Structures from the Protein Databank and Cambridge Structural Database. J. Chem. Inf. Model. 50, 572–584 (2010).

26. Computational approaches for protein complex modeling for intermediate resolution cryo-EM maps. 10.1016/bs.pmbts.2026.03.003 doi:10.1016/bs.pmbts.2026.03.003.

27. Wang, X., Terashi, G. & Kihara, D. CryoREAD: de novo structure modeling for nucleic acids in cryo-EM maps using deep learning. Nat Methods 10.1038/s41592-023-02032-5(2023) doi:10.1038/s41592-023-02032-5.

28. Zhang, Z. et al. Accurate Macromolecular Complex Modeling for Cryo-EM with CryoZeta. 10.64898/2026.02.13.705846(2026) doi:10.64898/2026.02.13.705846.

29. Vaswani, A., et al. Attention Is All You Need. Preprint at 10.48550/arXiv.1706.03762 (2017).

30. Hendrycks, D. & Gimpel, K. Gaussian Error Linear Units (GELUs). Preprint at 10.48550/arXiv.1606.08415 (2023).

31. Wu, Y. & He, K. Group Normalization. Preprint at 10.48550/arXiv.1803.08494 (2018).

32. Abramson, J. et al. Accurate structure prediction of biomolecular interactions with AlphaFold 3. Nature 630, 493–500 (2024).

33. Karras, T., Aittala, M., Aila, T. & Laine, S. Elucidating the Design Space of Diffusion-Based Generative Models. in Proceedings of the Advances in Neural Information Processing Systems 35: Annual Conference on Neural Information Processing Systems 2022 (2022).

34. Needleman, S. B. & Wunsch, C. D. A general method applicable to the search for similarities in the amino acid sequence of two proteins. Journal of Molecular Biology 48, 443–453 (1970).

35. Cragnolini, T., et al. *TEMPy* 2: a Python library with improved 3D electron microscopy density-fitting and validation workflows. Acta Crystallogr D Struct Biol 77, 41–47 (2021).

36. Zhang, C., Zhang, X., Freddolino, L. & Zhang, Y. BioLiP2: an updated structure database for biologically relevant ligand–protein interactions. Nucleic Acids Res 52, D404–D412 (2024).

37. Taniguchi, S., et al. ADOPT: Modified Adam Can Converge with Any $\beta_2$ with the Optimal Rate. in arXiv.org (2024).

